# Unbiased multiplex antigen screening of Cerebrospinal Fluid detects microbial and autoantigenic epitopes associated with Multiple Sclerosis

**DOI:** 10.1101/2024.07.05.602301

**Authors:** Nathaniel J. Barton, Khanh Tran, Meagan N. Olson, Mugdha Deshpande, Irina Radu, Nimmy Francis, Mariana Kurban, Adrian R. Orszulak, Samantha M. Chigas, Jonathan Sundstrom, Pepper Dawes, Liam F. Murray, Carolina Ionete, Elaine T. Lim, Christopher C. Hemond, Yingleong Chan

## Abstract

To comprehensively investigate the intrathecal antibody profile of multiple sclerosis (MS), we examined the cerebrospinal fluid of 195 patients (92 MS and 103 non-MS) for antibodies using a multiplex unbiased bacteria peptide library. We first tested against Epstein-Barr nuclear antigen 1 (EBNA1) for epitope sites enriched in MS and found a significant enrichment at position 407-419. We then used the data to perform a high-throughput screen against a library of 129 viruses known to infect humans. We discovered several additional epitopes from viruses such as Hantaan virus, Human Herpesvirus 6A and Human respiratory syncytial virus B associated with MS. Besides viral epitopes, we also screened for potential autoantigens of the central nervous system (CNS). We discovered several autoantigenic epitopes in proteins such as ADRB3, HTR3A and MPO that were significantly enriched for MS. Because of previous associations of Toxoplasma gondii infection with MS, we also performed a Toxoplasma gondii specific analysis and discovered additional epitopes enriched for MS. We further assessed epitope-epitope correlations within the patient samples and found distinct patterns of association between these microbial and autoantigenic epitopes. Finally, we performed machine-learning to determine if these epitopes are predictive for MS and found that the model incorporating all the epitopes could most effectively discriminate between MS and non-MS (ROC-AUC score = 0.91). Our results demonstrate the effectiveness of multiplex unbiased screens to detect the identity of potentially cross-reactive antibodies targeting MS CNS epitopes and they can also be used as effective biomarkers for MS.

**One Sentence Summary:** We performed an unbiased multiplex bacteria peptide antibody library screen on cerebrospinal fluid samples of patients with multiple sclerosis (MS) as well as non-MS controls and detected multiple viral and autoantigenic epitopes that are significantly enriched in MS patient samples.

## Introduction

Multiple sclerosis (MS) is an immune-mediated disease of the central nervous system (CNS), and the leading cause of non-traumatic neurological disability in young persons^1^. MS is primarily characterized by demyelination and injury of axons in the central nervous system (CNS), features that are frequently accompanied by neurodegeneration. The etiopathogenesis of MS remains incompletely understood; however, the role of antibodies has been suggested for decades, following the discovery of a polyclonal intrathecal antibody response (oligoclonal bands), with high specificity for the disease. Although no consistent antibody ‘target’ has emerged from these initial observations, the study of antibodies in MS has yielded numerous productive clinical discoveries in the past several decades^2^. These include the characterization of pure of antibody-mediated diseases formerly diagnosed as MS, including the aquaporin 4 antibody (now: NMO spectrum disease) and the Myelin Oligodendrocyte Glycoprotein antibody (now: MOG-associated disease)^3^.

Beyond just the potential to identify additional antibody-mediated disease, the study of intrathecal antibodies can also yield insight into the infectious history of patients with MS. Longstanding epidemiological evidence has suggested a role for infections in MS; most recently, strong evidence has emerged supporting a causative role for late infection with the Epstein Barr Virus (EBV)^4, 5^. MS is characterized by universal EBV seroprevalence^6^, with clinical disease manifestations following infection by an average of 10 years during which antibody titers directed against the EBV nuclear antigen (EBNA1) complex were found to have increased^7–10^. Notwithstanding this role for EBV, co-infections may also play an important role in the development or modulation of MS pathology during this period, with mixed support for a wide variety of microbes including bacteria (H. Pylori, Chlamydia pneumoniae, mycobacterium)^11–13^, parasites (Enterobius Vermicularis)^14^, and numerous viruses including torque teno^15, 16^, other herpesviruses (HSV-1, HHV-3, and HHV-6)^17–20^ and human endogenous retrovirus family (HERV)^21, 22^. Interestingly, the biomarker with the highest specificity for MS is the combination of antibody titers against the Measles, Rubella, and Varicella viruses (MRZ reaction) assessed from cerebrospinal fluid (CSF)^23, 24^.

One hypothesized mechanism of infectious-related injury in MS is the creation of potentially cross-reactive antibodies against CNS surface autoantigens. A recent study reported an autoantibody signature from a subset of patients with MS^25^. Another study showed that antibodies from serum and CSF MS samples bound more to the surface of oligodendroglial and neuronal cell-lines compared to control samples^26^. Also, previous studies have reported Myelin Proteolipid Protein (PLP), Myelin Basic Protein (MBP), myelin oligodendrocyte glycoprotein (MOG), myelin-associated antigen (MAG), myelin-associated oligodendrocyte basic protein (MOBP), and 2′,3′-cyclic-nucleotide 3′-phosphodiesterase (CNPase) as potential targets of these antibodies^27^. Besides these, there are also several studies reporting immune reactivity towards the EBV nuclear antigen protein EBNA1 through molecular mimicry/cross-reactivity with proteins of the CNS. A study demonstrated that the EBNA1 amino acid sequence 386-405 displayed pathogenetic mimicry/cross-reactivity with the CNS GlialCAM protein^28^. Other studies reported that residue 411-426 of EBNA1 cross-reacts with CNS myelin basic protein (MBP)^8, 29^, and residues 431-440^30, 31^ with the CNS potassium channel anoctamin 2 (ANO2). There is also evidence that alpha-crystallin B (CRYAB) is one of the targets of these cross-reactive antibodies^32^. These results suggest that once these cross-reactive antibodies hit their targets, they can trigger both direct cytotoxicity^33^ as well as perpetuation of ongoing inflammation through downstream recruitment and activation of immune cells including natural killer (NK) cells^34^. In light of such cumulative evidence, we aimed to better elucidate MS pathogenesis through comprehensive antibody epitope mapping of the CNS in MS patients.

Here, we report a study to characterize the global antibody epitope profile from the CSF of about 200 patients with and without MS. We utilized an unbiased randomized bacteria peptide library approach to map out the epitopes that are enriched in the CSF of MS versus non-MS patients^35, 36^. Because this assay uses a library of randomized sequences, it enables the ability to bioinformatically map the data onto any protein, as long as the protein’s amino-acid sequence is known. We performed such exploratory analyses on the EBV EBNA1 protein, a library of viral/microbial proteins as well as a library of potential human CNS autoantigens and reported our findings of epitopes detected to be enriched in MS patients compared to controls.

## Results

### The EBNA1 epitope at amino-acid position 408-419 is significantly associated with MS

We performed an unbiased multiplex screen using a random epitope library on CSF donor samples from 92 MS cases and 103 non-MS controls (**Table S1**). We tested the library for confounding effects due to age and sex, and we did not observe any significant correlations (**Figure S1**). The random epitope library allows us to analyze any potential protein for their significance for MS by mapping the data to the protein sequence (See Materials and Methods). As previous studies have consistently found a significant enrichment of antibodies targeting EBNA1 in MS blood sera and CSF^4, 5^, we initially analyzed our data from the multiplex screen by mapping the data to EBNA1. From this, we identified a significant epitope in EBNA1 from amino acids 407-419 (FDR = 5 x 10^-5^) to be associated with MS (**Figure 1A-C**). This result is consistent with previous studies, reporting the same epitope site being associated with MS^31,8,28^.

**Figure 1.**
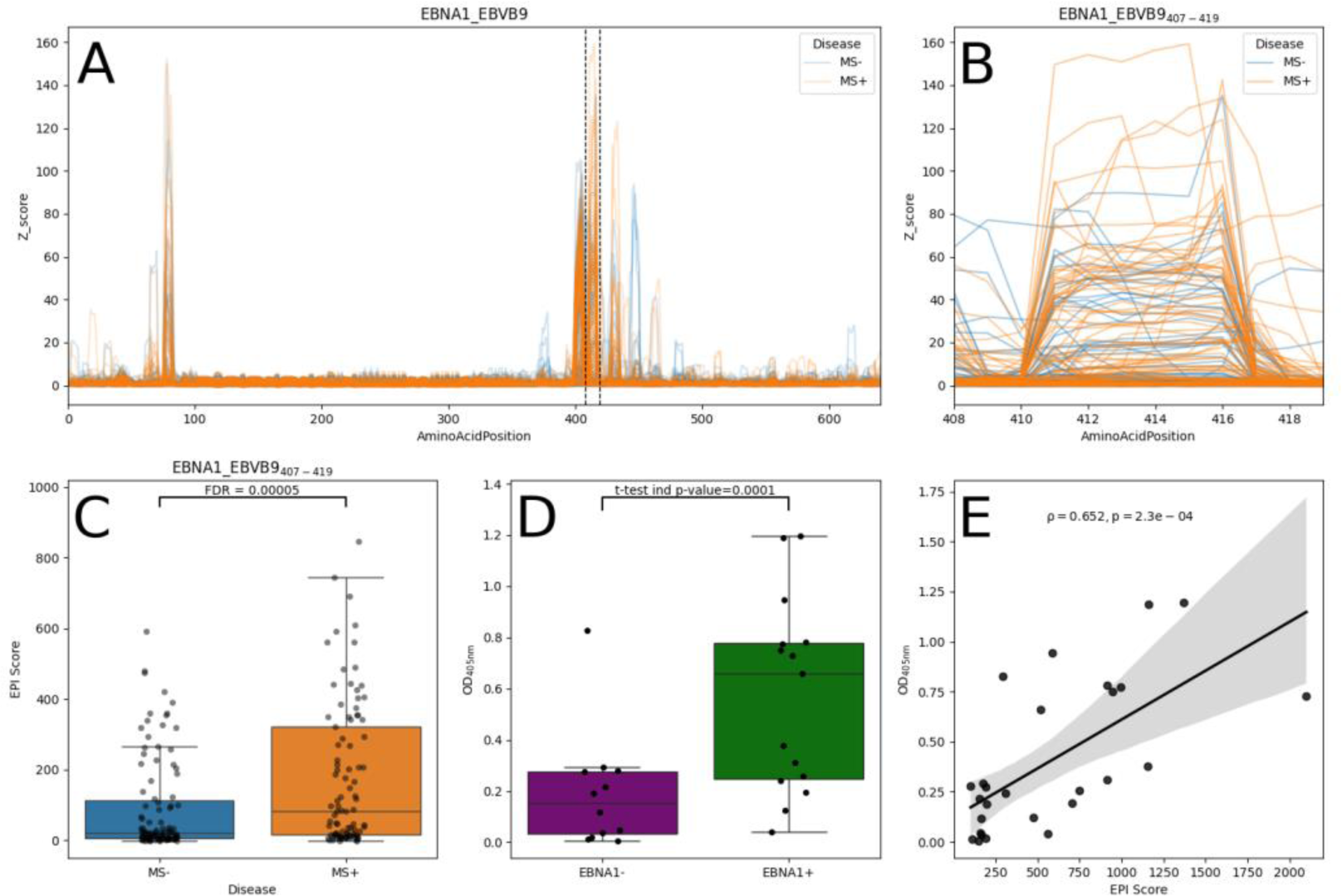
A) Line Plot of the epitope enrichment (expressed as a Z-score) for each amino acid position across the length of EBNA1 protein sequence. The peak (amino acids 407-419) is associated with MS. B) Zoomed-in version of A, showing the individual samples at amino acids 408-419. C) Boxplot showing the distribution of the epitope intensity score (EPI Score) for all samples at EBNA1_407-419_ with each point depicting an individual sample. MS cases are depicted as orange and non-MS controls are depicted as blue. D) Boxplot showing the distribution of OD 405 nm scores for ELISA results from CSF samples. EBNA1 positivity was predicted by the EBNA1 EPI Score. E) Scatterplot showing the correlation between the EBNA1 protein EPI Score and OD 405 nm ELISA results, with Pearson’s R correlation. For D and E, each point is the average of (n=2) replicates.

Besides the multiplex screen, we also independently verified the EBNA1 signal by enzyme-linked immunosorbent assays (ELISAs). We used the epitope intensity score (EPI Score), which is the cumulative epitope area under the curve across the EBNA1 protein and selected 27 CSF samples (based on sample availability) that would be either positive (EBNA1+) or negative (EBNA1-) for EBNA1 (**Table S2**) (See Materials and Methods). From our ELISAs, we observed a significant difference between EBNA1+ and EBNA1-samples (P = 1×10^-4^) (**Figure 1D**), which is indicative of the effectiveness of the multiplex antigen screen (**Figure S2**). Furthermore, there was a significant positive correlation between the EBNA1 EPI Score and the raw OD 405nm ELISA values (P = 7.5×10^-4^) (**Figure 1E**). Taken together, these results validate the predictive capability of the multiplex antigen screen.

### Screening across 129 viruses identified 22 epitopes significantly associated with MS

To investigate the role of infections with other viruses in MS besides EBV, we performed a screen of 3079 viral proteins from 129 human viruses to determine if there are epitopes from other viruses associated with MS (**Table S3**). From this analysis, we identified 18 viral epitopes that were significantly enriched for MS (FDR < 0.05) and an additional 4 were marginally significant (FDR < 0.06) (**Table 1**, **Figure 2**). These epitopes, which include EBNA1_407-419_, are contained within 17 different viruses with the epitope sequence ranging between 7 and 16 amino acids (**Table 1**, **Figure S3, Table S4**). Upon further examination, 3 of the significant epitopes were homologous to the ankyrin-repeat protein of pox viruses. 1 of the epitope was mapped to Cowpox (Uniprot ID: Q8QN44_CWPXB) and the other 2 were mapped to the same viral protein in different builds of Monkeypox (Uniprot ID: Q8UYF7_MONPZ and NCBI ID: YP_010377005.1). These epitopes share the consensus epitope sequence RRALIDEIIN (**Figure S4A**). Another two epitopes (Uniprot IDs: VB04_VAR67 and Q0GNP8_HSPV) shared the consensus epitope sequence VLNIN. This indicates that the signal from these epitopes are dependent and is a result of shared sequence homology.

**Figure 2.**
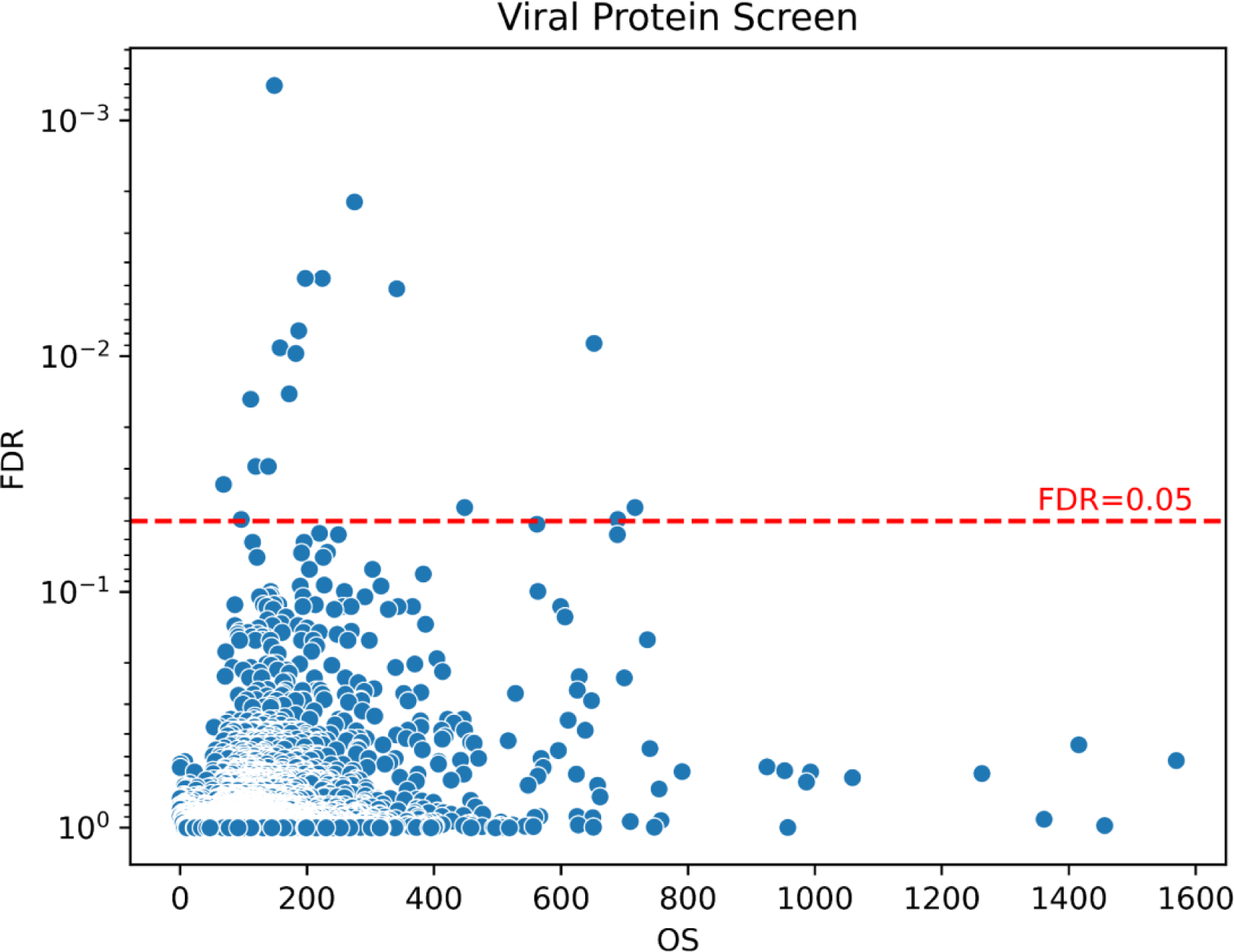
Screen of viral epitopes significantly associated with MS. Each point represents one viral epitope. 43387 epitopes were discovered across 3068 viral proteins. The epitope outlier sum score (OS) is represented on the x-axis. The Benjamini-Hochberg false discovery rate is represented on the y-axis. The red-dashed line represents FDR=0.05 and epitopes above this line are considered significantly associated with MS.

**Table 1.**
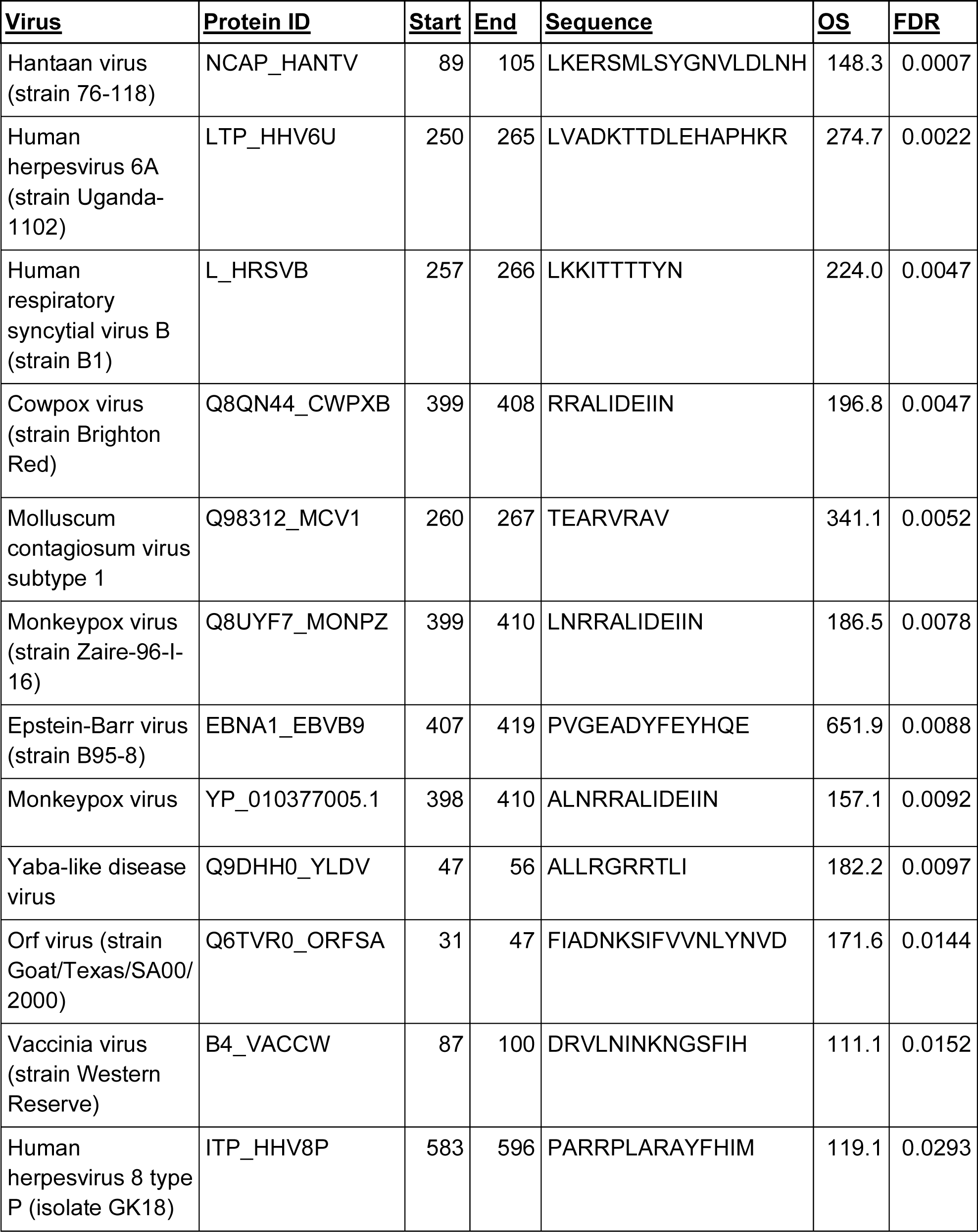
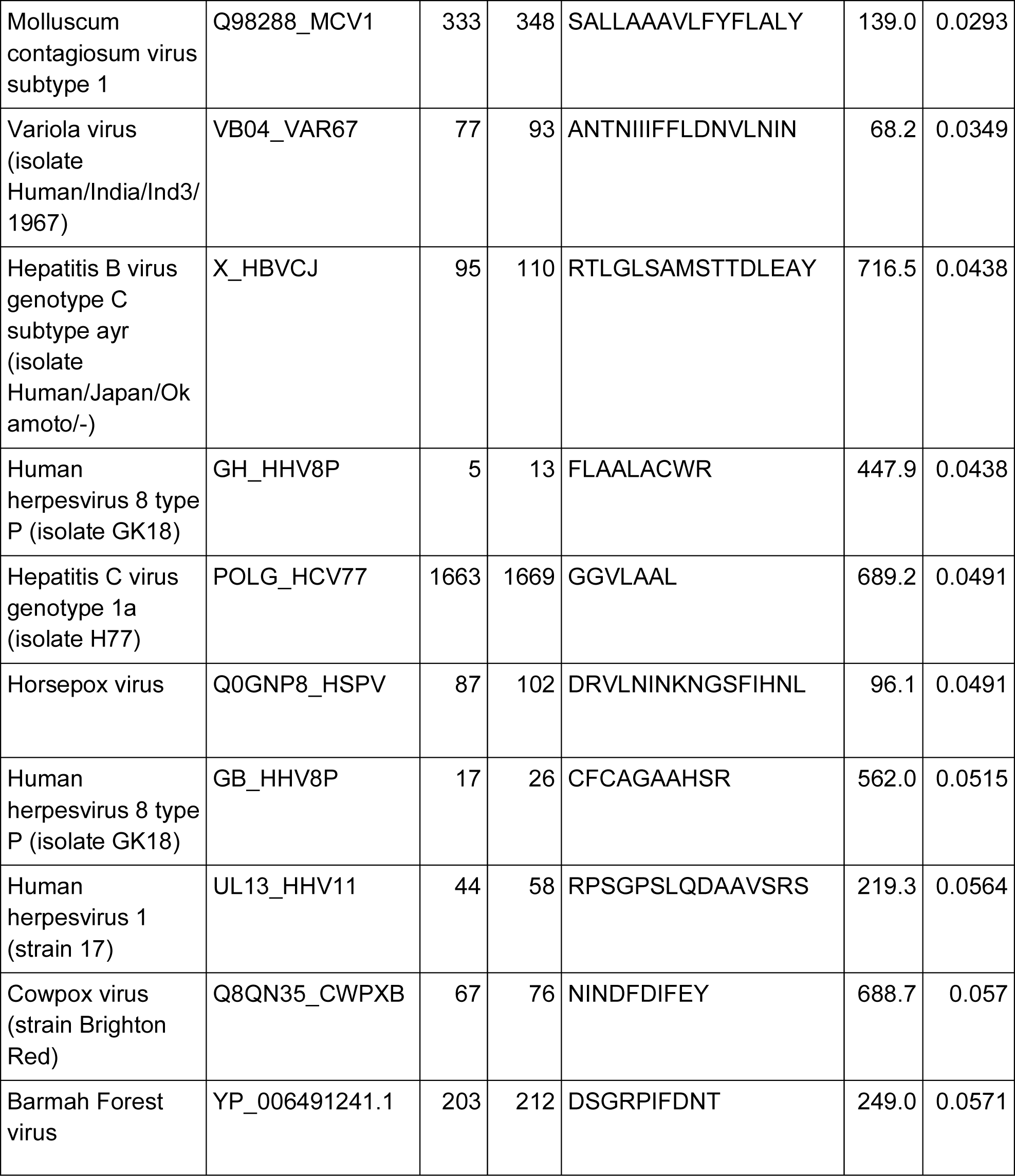
A list of significant viral epitopes detected from the unbiased screen. The columns are as follows: Virus–Name of the virus; Protein Name–UniProtKB/Swiss-Prot/NCBI ID of protein; Start– first amino acid position of the epitope; End– last amino acid position of the epitope; Sequence– amino acid sequence of the epitope; OS– the outlier sum score of the epitope; FDR– Benjamini-Hochberg false discovery rate.

We further investigated if any of the epitope sequences also mapped to proteins of other microbes by using the Basic Local Alignment Search Tool (BLAST)^37^. We detected that the epitope on Yaba-like disease virus (ALLRGRRTLI) shares sequence homology with proteins of other microbial species such as Spizellomyces Punctatus, Tanapox virus and multiple proteins of various Streptomyces bacteria. The epitope on Molluscum contagiosum virus (TEARVRAV) shares sequence homology with proteins on multiple other microbes such as Micromonospora pisi, Mesorhizobium and Podospora didyma. As such, while these epitopes align with the detected virus, their signal of association with MS may be due to infection from these other microbial species.

### Mapping the data to potential CNS autoantigens reveals several significant epitopes associated with MS

While the random epitope library was designed to detect viral epitope associations, we were also able to use the data to determine if potential autoantigens are associated with MS. There were several such autoantigens associated with MS and MS-related diseases from previous published reports. These include MOG^38, 8^, MAG^39^, ANO2^30, 31^, AQP4^39^, HEPACAM (GlialCAM)^28^, CRYAB^39, 32^, PLP^40, 41^ and MBP^42^. As an initial autoantigenic screen, we analyzed these proteins using our method to detect if there exist any epitopes in these proteins that are significantly enriched in MS samples. Our analysis identified 26 epitopes across these proteins, but we did not detect any epitope significantly associated with MS (FDR > 0.05) (**Table S5**). We also scored the specific epitopes identified in previous reported studies, but we were unable to detect any that were significantly associated (FDR<=0.05) with MS (**Figure S5, Table S6**).

Next, we expanded our analysis on a library of human proteins to detect for novel potential human autoantigens associated with MS. Rather than screening the entire human proteome, we curated a targeted list of 2438 human proteins known to be expressed on the surface of cells in the CNS (**Table S7**) (See Materials and Methods). Our screen identified 11 significant epitopes associated with MS (FDR < 0.05) and an additional epitope marginally associated with MS (FDR < 0.06) (**Figure 3**, **Table 2, Figure S6, Table S8**). Like with the viral epitopes, we performed a BLAST analysis and found that the ADRB3 epitope (PAPSRSLA) shares sequence homology with the RNA polymerase Rpb1 C-terminal repeat-containing protein of Toxoplasma gondii (**Figure S3B**). The DAGLB epitope (LGGGAAAL) shares sequence homology with several microbial proteins such as Myxococcales bacterium and Raphidocelis subcapitata. The remaining epitopes mapped to protein homologs of Mammalian or other Eukaryotic species.

**Figure 3.**
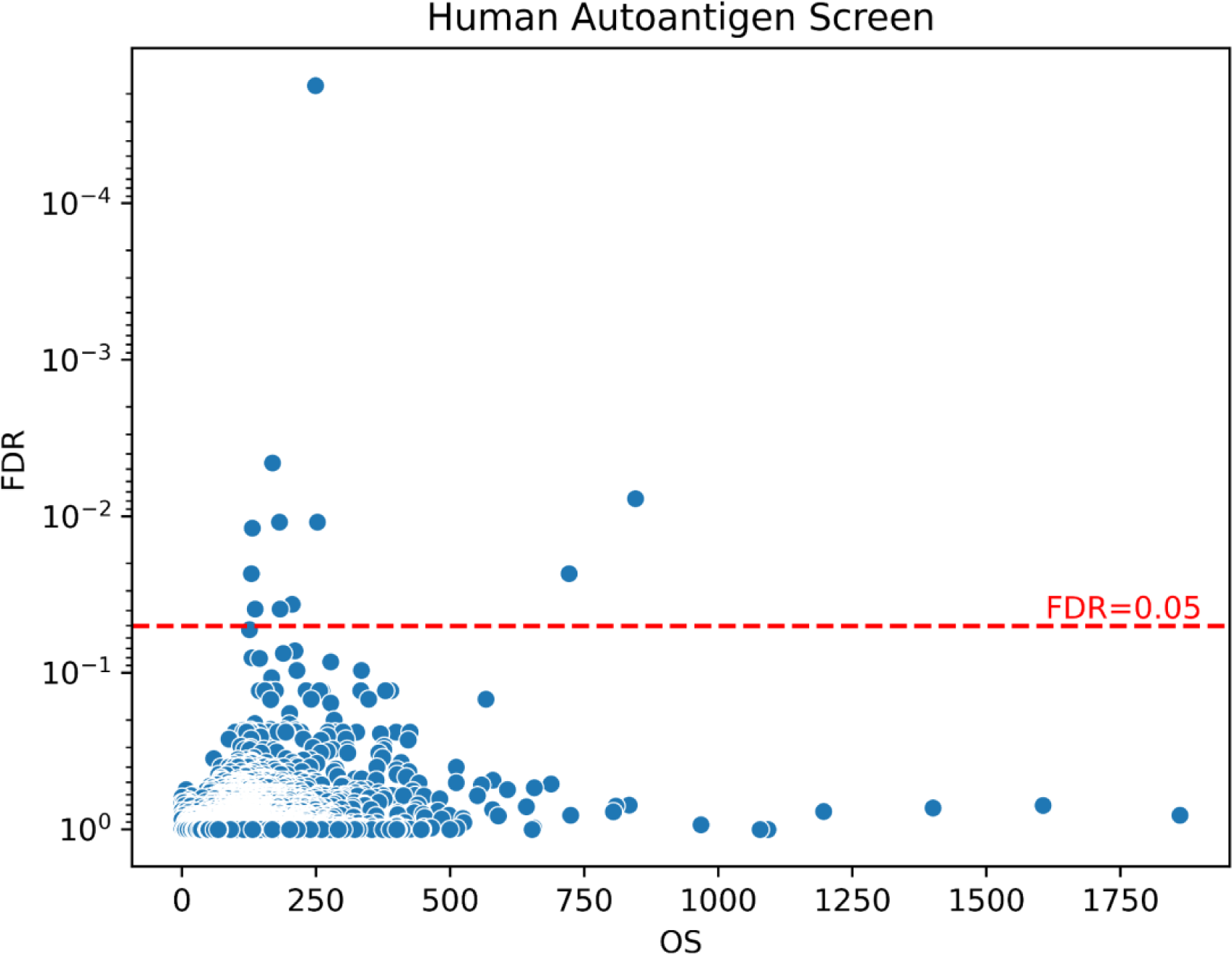
Screen of human autoantigens associated with MS. The epitope outlier sum score (OS) is represented on the x-axis. The Benjamini-Hochberg false discovery rate is represented on the y-axis. The red-dashed line represents FDR=0.05 and epitopes above this line are considered significantly associated with MS.

**Table 2.**
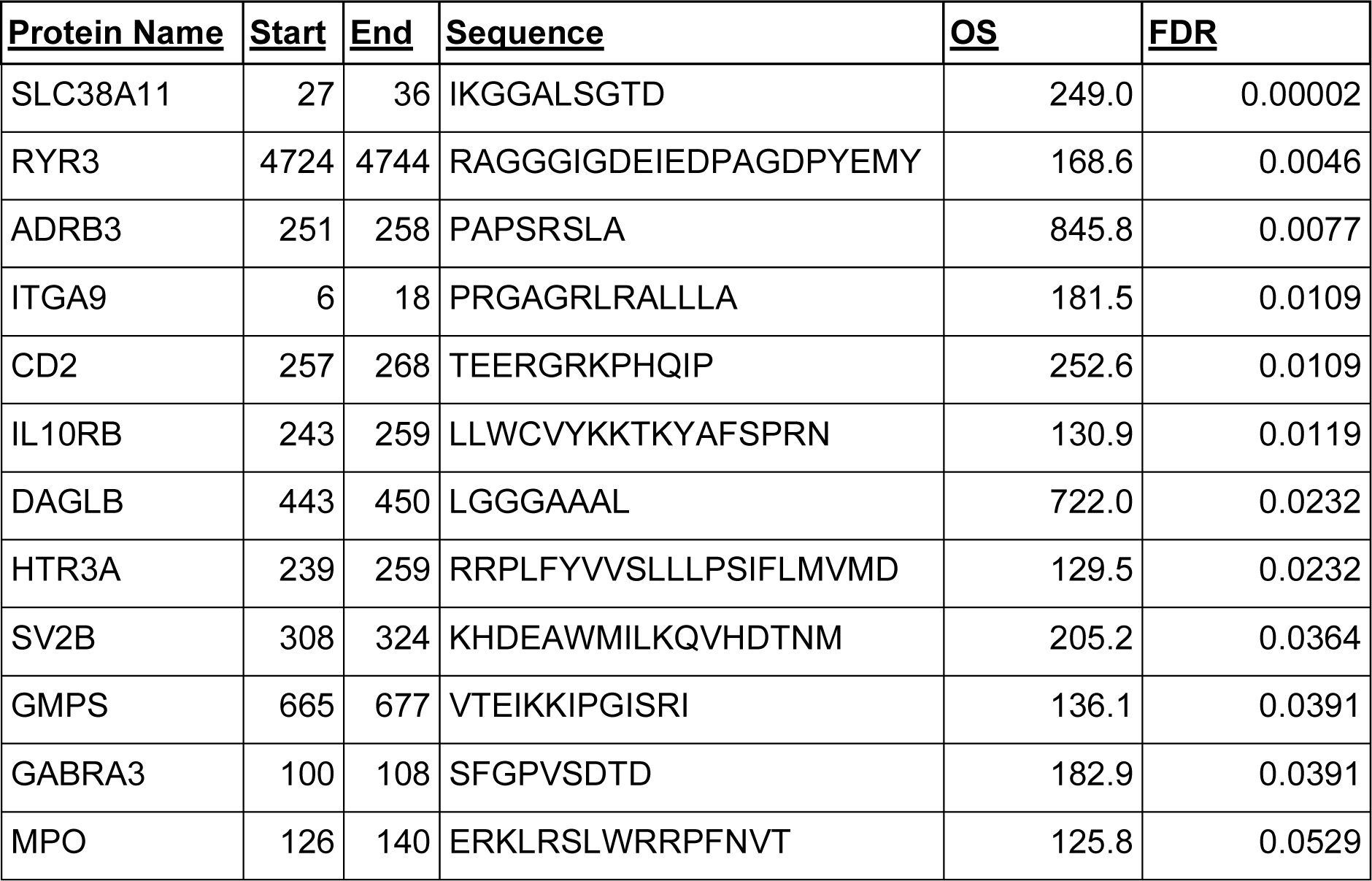
MS-associated human autoantigenic epitopes detected from the unbiased screen. Protein Name– HGNC protein symbol; Start-first amino acid position of epitope; End–last amino acid position of epitope; OS–Outlier sum score for epitope;FDR– Benjamini-Hochberg false discovery rate.

### Performing the screen on the Toxoplasma gondii proteome reveal significant associations of several epitopes with MS

Because we detected the ADRB3 epitope to be homologous with a Toxoplasma gondii protein and there are numerous reports linking Toxoplasma gondii with MS^43–48^, we decided to also map the data with the entirety of the Toxoplasma gondii proteome (**Table S9**). From this analysis, we identified 7 additional epitopes in the Toxoplasma gondii proteome that were significantly associated with MS (**Figure 4**, **Table 3, Figure S7, Table S10**). This result suggests that certain antigens on the Toxoplasma gondii proteome could generate cross-reactive antibodies that are potentially relevant for MS pathology.

**Figure 4.**
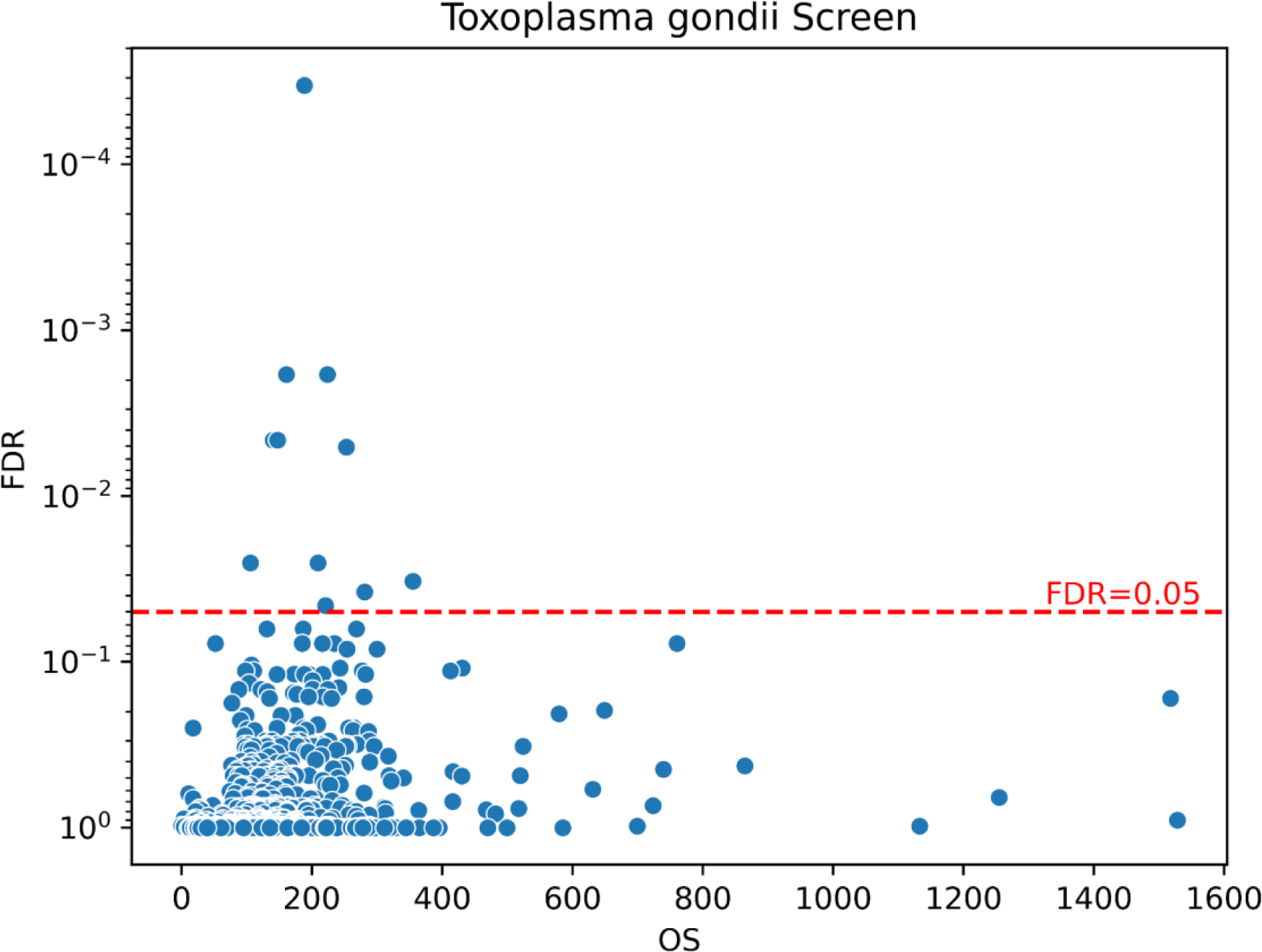
Screen of toxoplasma gondii epitopes associated with MS. The epitope outlier sum score (OS) is represented on the x-axis. The Benjamini-Hochberg false discovery rate is represented on the y-axis. The red-dashed line represents FDR=0.05 and epitopes above this line are considered significantly associated with MS.

**Table 3.** MS-associated Toxoplasma gondii epitopes. The columns are as follows: Protein Name–UniProtKB/Swiss-Prot entry name of protein; Start– first amino acid position of the epitope; End– last amino acid position of the epitope; Sequence– amino acid sequence of the epitope; OS– the outlier sum score of the epitope; FDR– Benjamini-Hochberg false discovery rate.

| **Protein Name** | **Start** | **End** | **Sequence** | **OS** | **FDR** |
| --- | --- | --- | --- | --- | --- |
| Q1JTC7_TOXGO | 149 | 168 | ETAGDNGSRSRFEKKNEKLG | 188.4 | 0.00003 |
| Q1JTB7_TOXGO | 457 | 467 | FEEFEHATGGC | 161.0 | 0.0018 |
| Q1JSA6_TOXGO | 373 | 392 | VRAAMQAQVAQYLEADRRP<br>R | 223.8 | 0.0018 |
| Q1JSL8_TOXGO | 159 | 174 | EWGACEGEARPGRGPL | 140.6 | 0.0046 |
| Q1JSS8_TOXGO | 450 | 469 | EDEGEAGTDEVSDTPPERIF | 147.2 | 0.0046 |
| Q1JTI9_TOXGO | 257 | 267 | VVEAQARRAVS | 252.7 | 0.0051 |
| Q1JT23_TOXGO | 1312 | 1326 | AVSRARFLLEEIRHQ | 209.2 | 0.0253 |
| Q1JTK6_TOXGO | 76 | 85 | QRRPLGLKRG | 105.6 | 0.0253 |
| Q1JTJ5_TOXGO | 323 | 335 | QTVPAANSAEAYV | 354.9 | 0.0327 |
| Q1JSR1_TOXGO | 908 | 916 | RETALAHHG | 280.9 | 0.0379 |
| Q1JSX1_TOXGO | 650 | 663 | EGQEAHAEQKERGT | 221.0 | 0.0458 |

### Specific viral and Toxoplasma gondii epitopes correlate with human autogenic epitopes

To investigate whether viral or microbial infections can result in cross-reactive autoantibodies targeting CNS autoantigens resulting in MS, we looked at the EPI Score correlations between the detected epitopes associated with MS (**Figure 5A**). While we found many correlations between epitopes not specific to MS, some correlations stood out as being meaningful. We observed strong correlations between several viral and toxoplasma gondii epitopes with the ADRB3, HTR3A and the MPO autoantigenic epitopes suggesting that these microbial infections could result in the generation of antibodies that target these autoantigens resulting in increased MS risk (**Figure 5A**).

**Figure 5.**
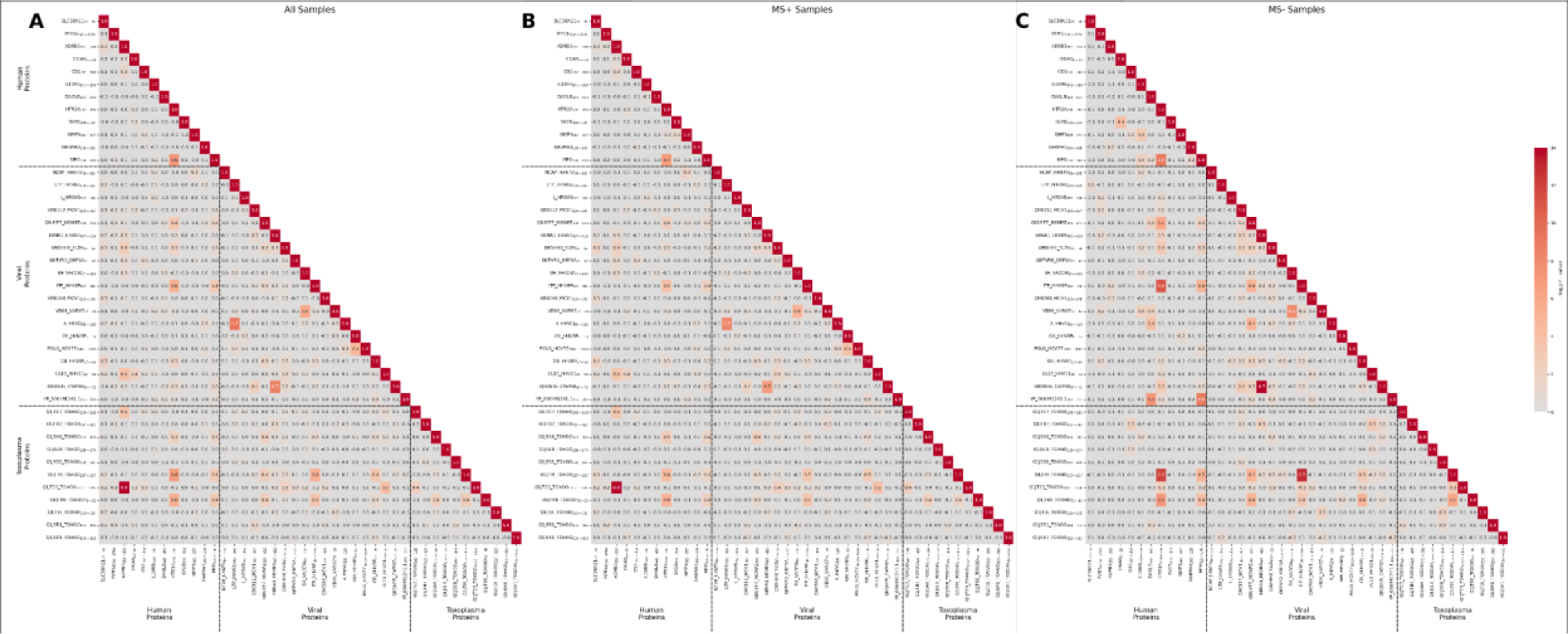
Heatmap of the pairwise correlations of the epitope intensity score (EPI Score) between all the epitopes for (A) All samples, (B) MS only samples and (C) Non-MS samples. The value of each cell is the Pearson’s correlation coefficient of the pairwise epitope comparison while the color of the cell depicts the -log_10_ P-value of the association.

To investigate whether any of these correlations are MS-specific, we separately analyzed the correlations only with MS patient samples as well as only with non-MS patient samples (**Figure 5B-C**). First, we find that a Toxoplasma gondii epitope (Uniprot ID: Q1JT23_TOXGO, amino-acid position 1312 to 1326) and several other viral epitopes have a strong MS-specific correlation with the ADRB3 epitope (**Figure 6A**). While this suggests that Toxoplasma gondii infection can generate autoantibodies against ADRB3 to cause MS, it could also be explained by Toxoplasma gondii infection generating antibodies against Toxoplasma gondii itself, as our earlier result shows that the ADRB3 epitope shares sequence homology with a Toxoplasma gondii protein. Next, we observed a strong MS-specific correlation between the epitope on Human herpesvirus 6A (Uniprot ID: LTP_HHV6U) and the epitope on the Hepatitis B virus (Uniprot ID: X_HBVCJ) suggesting that co-exposure to HHV6A and HBV due to infection or vaccination is strongly linked with MS (**Figure 6B**). We also observed a similar pattern between the epitope on Hepatitis C virus (Uniprot ID: POLG_HCV77) and Human herpesvirus 8 (Uniprot ID: GB_HHV8P) (**Figure 6C**).

**Figure 6.**
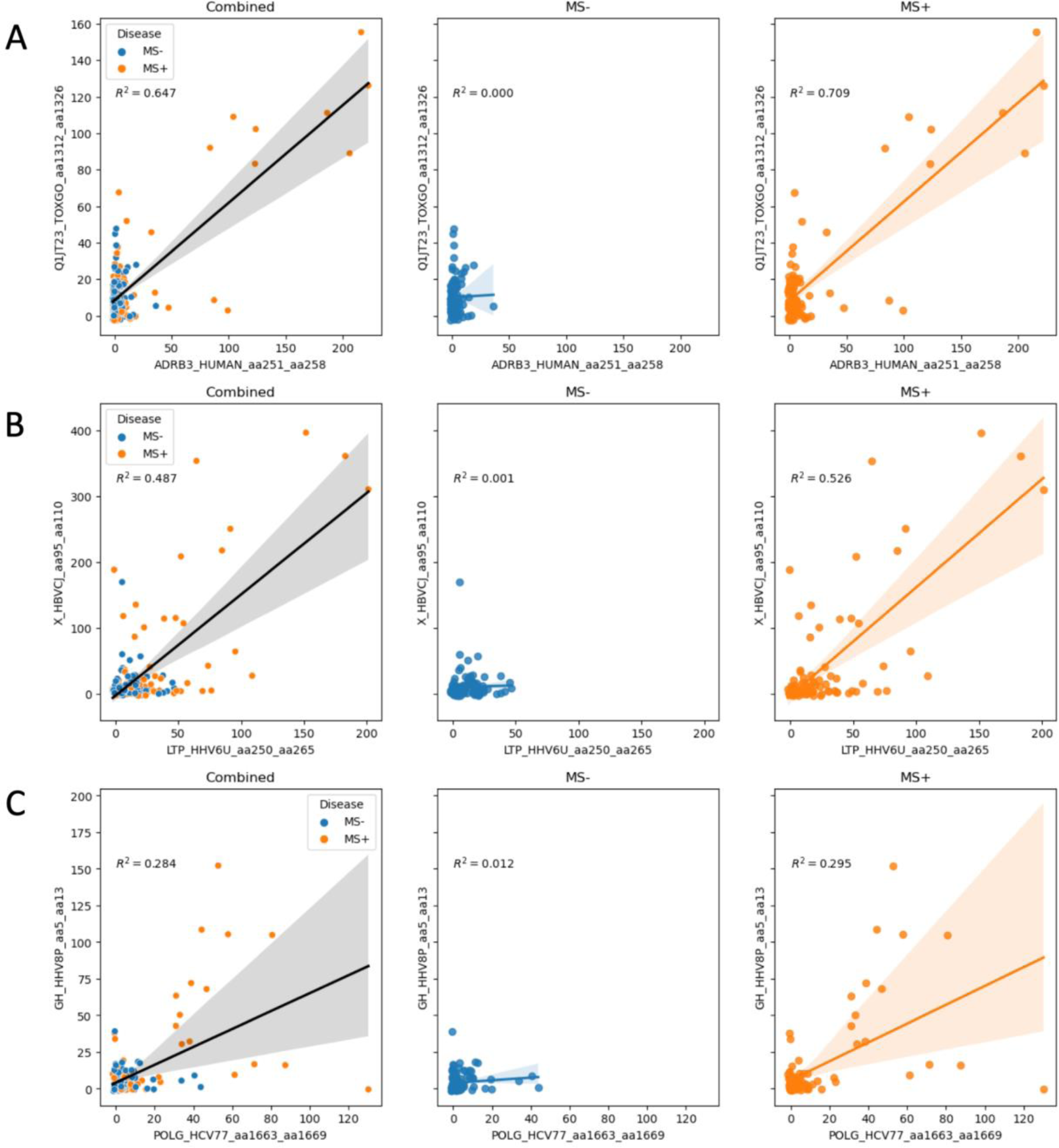
Correlation plots for specific epitope pairs that show MS-specific correlation. (A) Plot between the ADRB3 epitope and the Toxoplasma gondii epitope (Q1JT23). (B) Plot between the human herpesvirus 6 epitope (LTP_HHV6U) and the hepatitis B virus epitope (X_HBVCJ). (C) Plot between the hepatitis C virus epitope (POLG_HCV77) and human herpesvirus 8 epitope (GH_HHV8P). The axes represent the corresponding epitope intensity score (EPI Score) for each sample. The leftmost plots show a combined plot for all samples whereas the middle and rightmost plots show non-MS correlations and MS-specific correlations respectively.

### A machine-learning model from the discovered epitopes can accurately predict MS

After identifying 42 non-overlapping unique epitopes across human, viral, and toxoplasma gondii proteins, we next asked if these epitopes could be used to predict MS status in patients given the epitope profiles. To answer this question, we developed four support vector machine (SVM) models, with cross validation, trained on 1) the 12 human epitopes, 2) the 19 viral epitopes, 3) the 11 Toxoplasma gondii epitopes, and 4) a combination of all 42 epitopes (See Materials and Methods). From our results, we find that all models were on average more than 60% accurate (**Figure 7A**) and the combined epitope model was much more accurate (∼ >80% probability of classifying MS versus non-MS). Furthermore, receiver operator characteristic analysis shows that the combined model could most effectively discriminate between true and false positives for MS, with a ROC-AUC score of 0.91, higher than any individual screen model (**Figure 7B**). Taken together, our machine learning model shows that these epitopes can be used to predict MS, and could be used as clinical biomarkers for MS.

**Figure 7.**
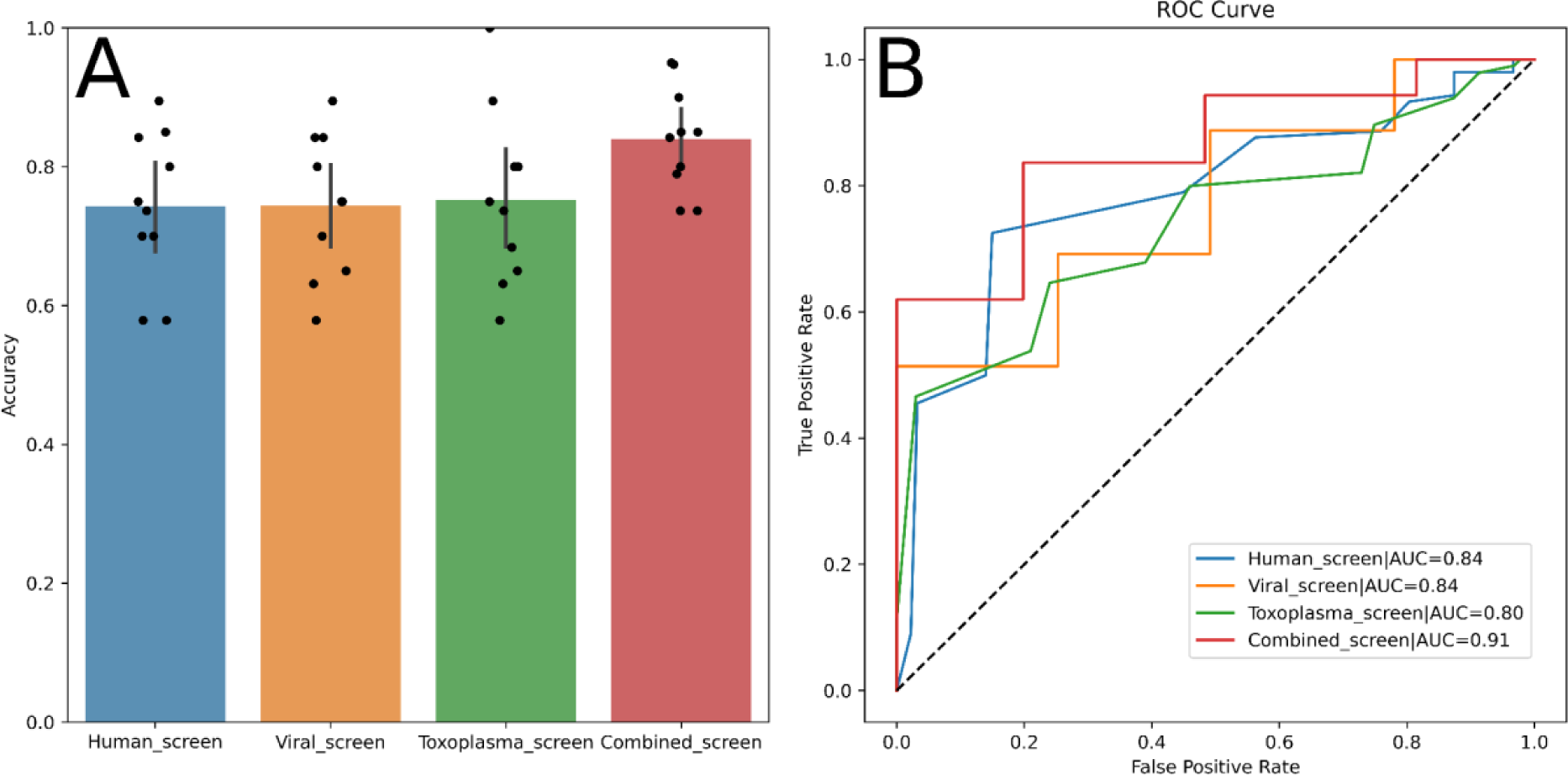
Machine learning model for predicting MS. A) Barplot of the accuracy of each SVM model across 10 cross-validation iterations. Error bars represent the 95% CI. B) Receiver Operating Characteristic (ROC) curve analysis comparing SVM models trained on either human epitopes (blue line), viral epitopes (orange line), Toxoplasma gondii epitopes (green line), or a combination of all three epitopes (red line). The black dashed line shows the expected result of a random classifier.

## Discussion

In this study, we describe the use of an unbiased multiplex bacteria peptide library combined with bioinformatics analyses for identifying epitopes associated with MS in patient CSF samples. Because we assay the antibodies within CSF samples, our data measures the antibody content within the CNS instead of the periphery. We discovered 42 non-overlapping unique epitopes within viruses, the toxoplasma gondii proteome as well as human autoantigens of the CNS significantly associated with MS. These epitopes reveal specific patterns of correlation between predicted viral, toxoplasma gondii and human autoantigenic epitopes and collectively, they can be accurately predictive for MS using machine learning models.

Our results show the potential relevance of other viruses besides EBV on MS pathology. First we detected several epitopes within viral proteins of the Poxviridae family of viruses such as Variola, Vaccinia, Coxpox, etc. Early researchers have suggested that pox viruses can be involved in MS because there are viral proteins with multiple domains that are like human host proteins such as GlialCAM, tenascin, neuregulin and contactin^49^. However, the viral MS epitopes that we detected were not in regions of homology in these human proteins suggesting other mechanisms for their association with MS. Besides pox viruses, there is a wealth of previous research linking human herpesvirus 6 (HHV-6) serostatus with MS in multiple studies of patient samples^50,20,51^. In our results, we detected an epitope on the Large tegument protein deneddylase of HHV-6 to be associated with MS. Besides HHV-6, we also discovered protein X of hepatitis B virus (HBV) to be associated with MS. HBV vaccination has been implicated to be associated with MS and some studies have suggested an increase in risk while others suggested no effect or a protective effect against MS^52–56^. Our results suggest that there is an association between HHV-6 and HBV as we observed a strong MS-specific correlation between their epitope signal. Besides HHV-6 and HBV, we also discovered epitopes on HSV-1 associated with MS. Previous reports also suggest that HSV-1 infection plays a role in MS^57^. Our results implicate the epitope region on UL13 being associated with MS. Our results suggest that these viral epitopes, in conjunction with EBV, can explain MS incidence better than EBV alone.

The results from the toxoplasma gondii analysis suggest that toxoplasma gondii infection is associated with MS^44, 48^. We perform this analysis because the detected ADRB3 epitope shared sequence homology with a toxoplasma gondii protein. These observations seem to contradict some earlier reports of toxoplasma gondii being protective against MS^43, 46^. There were also previous studies that did not find any association or even positive association with MS^58, 48^. One possibility would be that certain epitopes of toxoplasma gondii do cause risk due to antibody cross-reactivity but the general population do not develop these cross-reactive antibodies from toxoplasma gondii infection. We speculate a genetic or environmental influence on whether the development of anti-Toxoplasma gondii antibodies may be pathogenic, neutral, or protective for MS, which is a future research direction of ours.

Because of previous publications indicating molecular mimicry of cross-reactive antibodies linking EBV with MS^30,31,28,32^, we also screened for potential autoantigens by testing a library of CNS surface antigens. While we did not observe any of the previously reported epitopes, the results from the autoantigenic screen yield several interesting targets. We discovered an epitope site on HTR3A, a serotonin receptor associated with MS. The HTR3A epitope signal is also strongly correlated with several viral and toxoplasma gondii epitopes in both MS and non-MS samples. From previous studies, there are multiple lines of evidence to suggest that the serotonin pathway is protective of MS through the reduction of inflammatory cytokines such as IFN-y and IL-17^59^. In experimental autoimmune encephalomyelitis (EAE) mice, a mouse model of MS, it was reported that EAE mice had lower serotonin content compared to wildtype mice^60^. As such, this association with MS can be explained by cross-reactive antibodies acting as HTR3A antagonists. From our data, we also discovered an epitope on myeloperoxidase (MPO) associated with MS. This epitope is also highly correlated with the HTR3A epitope. MPO is a toxic enzyme promoting inflammation and its expression has been previously associated with MS^61, 62^. Our results indicate that CSF antibodies against MPO are increased in MS patients, which is consistent with previous literature.

Genes of some other significant autoantigenic epitopes were also previously associated with MS. There is evidence showing that Solute carrier family 38 member 11 (SLC38A11) is differentially expressed in the B-cells of MS patients compared to controls^63, 64^. Another associated gene is Integrin-alpha9 (ITGA9). ITGA9 forms a heterodimer with integrin-B1 and there are several MS related immune consequences reported about ITGA9^65^. In EAE mice, ITGA9 regulates the secretion of S1P receptors, which affect discharge of immune cells via the MAPK pathway^65^. Also, XCL1 serum levels are significantly higher in MS patients compared to healthy control. XCL1 is a ligand for ITGA9, and antibodies neutralizing XCL1 result in abolished disease progression in EAE mice^66^. There are also reports about interleukin 10 (IL10) involvement with MS. While interleukin 10 receptor subunit beta (IL10RB) has not been specifically implicated in MS, it was reported that IL10 serum levels are lower in MS patients compared to healthy controls^67^. It was also reported that EBV encodes its own IL10 homolog with high amino-acid identity with human IL10^68^. A study also reported that regulatory B-cells secreting IL10 are restored to normal levels after B-cell depletion therapy for MS^69^. The association of GABRA3 has also a prior association with MS. An early study showed statistically significant association of GABRA3 alleles with MS^70^. GABRA3 encodes for a GABA receptor containing an alpha 3 subunit, which is a receptor of the GABA family that responds to the GABA neurotransmitter. Several previous studies have demonstrated a connection between GABA function and MS. A previous study reported that inflammation in MS that resulted in altered cytokine levels within the CSF is associated with attenuation of GABAergic activity^71^. A more recent study demonstrated increased GABAergic receptor expression in MS patients in-vivo^72^. Another study also demonstrated a similar increase of GABAergic receptor density in MS^73^. Collectively, these reports demonstrate the potential biological mechanisms of these epitope’s involvement with MS pathology.

While we have discovered a comprehensive list of epitopes associated with MS, we also note that these epitopes were discovered using a random epitope library aligned to a list of known protein sequences. The analysis does not account for genotype variation resulting in coding variants within donor samples which could affect the results. There is also homology between different proteins and while we predict that the detected signals are the result of antibodies targeting our target proteins, the signals could arise from other proteins with sequence similarity. Also, the bacteria library generates surface peptides without post-translational modifications (PTMs) and any antibody targeting proteins specifically with PTMs will not be detected by the bacteria peptide library.

Our unbiased antibody library screening approach demonstrates its ability to detect antibodies against a wide variety of targets associated with MS. The advantage of using an unbiased random multiplex system is that the data generated can be used for detecting associated epitopes for any protein of interest with the help of innovative bioinformatics. We have detected viral, toxoplasma gondii as well as autoantigenic epitopes strongly associated with MS. We also demonstrated that these epitopes together have high probability for predicting MS from patient CSF samples and can be used as biomarkers for MS. The data reported here will provide researchers new avenues for understanding MS pathology as well as uncover biomarkers as well as novel therapeutic targets for treating MS.

## Materials and Methods

### Cohort description

#### Sample Collection

Cerebrospinal fluid (CSF) samples were obtained as part of routine clinical care from a single-center, prospective observational study (NeuroInflammatory Chronic Illness Study) at the University of Massachusetts Memorial Medical Center and Chan School of Medicine. As part of this study, patients may opt to donate a small portion of their CSF for research purposes after providing written informed consent. CSF samples are processed within 1 hour from the time of the lumbar puncture, centrifuged at 300G for 5 minutes, and aliquots created from the supernatant that were frozen at −80° C. None of the samples in this study were previously thawed. Patient clinical details were obtained from chart review of the electronic medical record.

#### Sample Description

We obtained a total of 210 CSF samples from patients. There were a total of 92 unique MS cases, 103 unique non-MS controls and 11 unique donor samples that had MS clinical and/or MRI features but did not meet diagnostic criteria for MS (**Table S1**). The data generated from these 11 samples were used in the development of the pipeline for epitope discovery but they were excluded when testing if the epitopes are statistically associated with MS. Also, there were 2 MS donors and 2 non-MS donors that had duplicate samples (**Table S1**). Raw data from these duplicated samples were generated independently, but the processed scores were averaged between the duplicates for detecting epitopes.

### Detecting and Testing epitopes for association with MS

#### Obtaining 5 and 6-mers Enrichment Scores

We acquired data for the random 12-mer bacteria epitope library by sending the CSF samples to Serimmune (serimmune.com) and running them through their Serum Epitope Repertoire Analysis (SERA) Platform. Briefly, they processed the samples by screening for IgG antibodies that bind to the 12-mer peptide antigen expressed on the E.coli bacteria library. The selected bacteria library was then lysed and sequenced by next-generation sequencing and read-counts for each 12-mer peptide was generated with approximately 3 million reads per sample.

Normalized k-mer enrichment scores for each sample were calculated for each k-mer based on the number of 12-mer reads, as previously described in the PIWAS method^36^. The enrichment score is calculated by first decomposing the sequenced 12-mers in constituent k-mers (with k=5 and k=6). For each k-mer, the enrichment score (E_s_) for each sample (S) is calculated as:

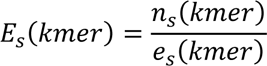

where n_s_(k-mer) is the number of unique 12-mers containing the k-mer and e_s_(kmer) is the the expected number of k-mer reads based on the amino acid proportions, defined as:

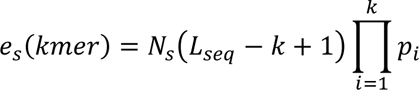

where N_s_ is the number of 12-mer reads generated for S, L_seq_ is the length of the amino acid reads (12 bases), k is the k-mer length, and p_i_ is the amino acid proportion for the i-th amino acid in kmer in all 12mer reads from S.

### Testing peptide counts for sex and age specific effects

For each sample, we sum the read count of all unique 12-mers and obtained an overall peptide count for each sample. We performed a Pearson correlation test between the age of individuals and their corresponding peptide counts using the pearsonr function from the SciPy library in Python to evaluate age-specific effects. An independent t-test was conducted to compare peptide counts between male and female samples using the ttest_ind function from the SciPy library in Python to evaluate sex-specific effects. All statistical analyses were performed using Python’s SciPy library. Significance levels were set at α = 0.05.

### Protein Tiling and Permutation

For a given amino acid sequence form a protein of interest (P), we next tiled the 5-mers or 6-mers (k) across the length of the protein and calculated the sum (T) of all overlapping enrichment scores at that position(i):

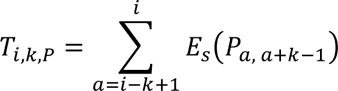

where P_a,a+k-1_ is the sequence of the k-mer found at position a to position a+k-1 on the protein (P).

Next, we permuted the amino acid sequence in 12-amino acid wide windows across the entire length of the protein and retiled the permuted area to get a null distribution (R):

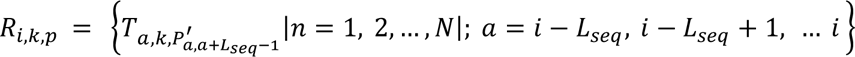

where N is the number of permutations (1000) and 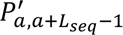 is the sequence of protein (P) with the amino acids in position a to a+L_seq_-1 randomly shuffled.

Using R_i,k,p_ as a null distribution, we calculate the Z-score for each amino acid as:

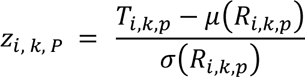

where μ and σ are the mean and standard deviation of R_i,k,p_ respectively.

Finally, we take the highest Z-score of each amino acid as:

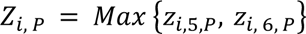

For donors with duplicate samples, the donor Z-score was obtained by averaging the Z-score (Z_i,P_) of all samples.

### Epitope Identification

Epitopes were identified via peak calling. For a given sample/protein, we plotted the amino acid position along the x-axis and the Z-score along the y-axis. We defined an epitope as a local maxima with a height of at least 7 (reflective of the 97.5th percentile of peak heights) (**Figure S8**) and a width of 6-15 amino acids (reflecting the typical epitope size recognized by antibodies). Peak calling was performed using Scipy’s signal.find_peaks function (https://scipy.org/citing-scipy/).

After identifying peaks across all samples, we next clustered peaks into epitopes. We defined a cluster as peaks within 20 amino acids of each other. For a cluster to be considered a distinct epitope, we required the peak to be found in at least 5 of the samples. Clustering was performed using complete-linkage clustering as implemented in Scipy’s cluster.hierarchy.fclusterdata function.

### Epitope Scoring

To compare epitopes, we calculated the area under the curve for each peak in the cluster from the peak’s left (l) to right (r) interpolated position, as calculated by Scipy’s signal.find_peaks function for the protein (P). We term this quantitative area as the epitope intensity score (EPI Score). For samples negative for a significant peak, we instead calculated the EPI Score from the leftmost peak (l) to the rightmost peak (r) in the set of samples positive for that peak. EPI Scores were calculated using the trapezoidal rule, as implemented in by the python sklearn module.

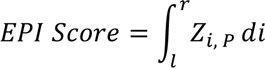

### Outlier Sum Calculation

For each epitope, we calculated the outlier sum statistic (OS) as described by Tibshirani and Hastie^74^. The outlier sum is more powerful than the usual t-statistic at finding differences in case/control groups where differential expression occurs in only a subset of cases. For each epitope (i) and sample (j), the EPI Score is normalized (*x^′^_ij_*) by dividing the difference of the EPI Score and the median EPI Score (med_i_) in control samples by the median absolute deviation in control samples (mad_j_) (in instances where the median absolute deviation was 0, we forewent dividing the difference) as follows:

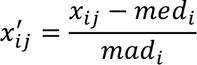

Next, the interquartile range (IQR) is calculated as the difference between the 75th and 25th percentile EPI Score of all samples (case and control) and values in the case set greater than the 75th percentile + IQR of all samples are considered outliers. All case outliers are then summed together to get the outlier sum statistic, as follows:

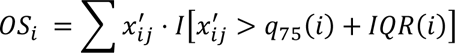

After calculating OS for each sample, we perform 100,000 permutations of the case/control labels to use as the null distribution to calculate a Z-score and p-value. P-values were corrected for multiple hypothesis correction using the Benjamini-Hochberg method.

### EBNA1 IgG ELISA Assay

We performed enzyme linked immunosorbent assay (ELISA) to detect EBNA1 human IgG antibodies (SERION ESR1363G) in 6 MS+ EBNA1+ samples, 5 MS+ EBNA1-samples, 9 MS-EBNA1+ samples, and 7 MS-EBNA1-samples (**Table S2**). EBNA1 positivity was determined based on their EPI Scores for EBNA1. CSF from MS patients and controls underwent dilution with a dilution buffer (SERION B231-S1), resulting in ratios of 1:4 and 1:2, respectively. Subsequently, these diluted samples were added to a 96-well plate coated with EBNA1 antigen. Each sample’s ELISA was conducted in a single replicate with a 1:2 ratio and in duplicates with a 1:4 ratio. The ELISA procedure adhered to the manufacturer’s instructions, and the readout, measured at an absorbance of 405 nm, was obtained using a plate reader (Perkin Elmer Victor X5) (**Figure S2**).

### Viral Screen

We obtained a list of 129 viruses that infect humans from the ViralZone project.^75^ Proteomes for each virus were then downloaded from Uniprot.^76^ For each protein, epitopes were identified and scored as explained above. FDR correction was applied to the list of all identified epitopes across all viruses.

### Human CNS Autoantigen Screen

We curated a list of potential autoantigenic target proteins in the central nervous system (CNS) by identifying proteins expressed on cell surfaces throughout the CNS. We combined a list of known and predicted surface proteins identified by the *in silico* human surfaceome project^77^ with a list of brain proteins from the Human Protein Atlas project ^78, 79^ to create a list of 2438 potential autoantigenic proteins in the CNS. Protein sequences for each protein were obtained from Uniprot^76^ and epitopes were identified and scored as explained above. FDR correction was applied to the list of all identified epitopes across all human proteins.

### Toxoplasma gondii Screen

We obtained the complete proteome for Toxoplasma gondii from Uniprot^76^ (Uniprot Proteome ID: UP000002437). The proteome contained 466 genes and epitopes were identified and scored as explained above. FDR correction was applied to the list of all identified epitopes across all Toxoplasma gondii proteins.

### MS prediction from epitopes

For each group of epitopes, a support vector machine (SVM) model was trained using the EPI scores calculated for each epitope. EPI scores from MS and non-MS patients were first standardized for each epitope. Training data was split using k-Fold cross-validation with k=10. SVM models were implemented using the scikit-learn SVC model^80^. The model outputs a prediction score ranging from 0 to 1. Scores >0.5 were classified as MS+ and scores ≤0.5 were classified as MS-for the calculation of accuracy scores. Accuracy scores were calculated by taking the number of correct predictions divided by the total number of predications. Area under the ROC Curve (ROC-AUC) and accuracy scores were also calculated using scikit-learn.

## Acknowledgements

We thank the patients who generously contributed their samples in this study for research. The use of patient CSF samples for research in this work was approved by the IRB of UMass Chan Medical School (Neuroinflammatory Chronic Illnesses, Study ID: 14143). This work was partially funded by the Dan and Diane Riccio Fund for Neuroscience (7/1/22 – 6/30/23).

## Data Availability

The data presented here will be made available upon request by contacting the corresponding authors.

## Declarations

None of the authors have any competing interests.

