## Supplemental Figures for "Unbiased multiplex antigen screening of Cerebrospinal Fluid detects microbial and autoantigenic epitopes associated with Multiple Sclerosis"

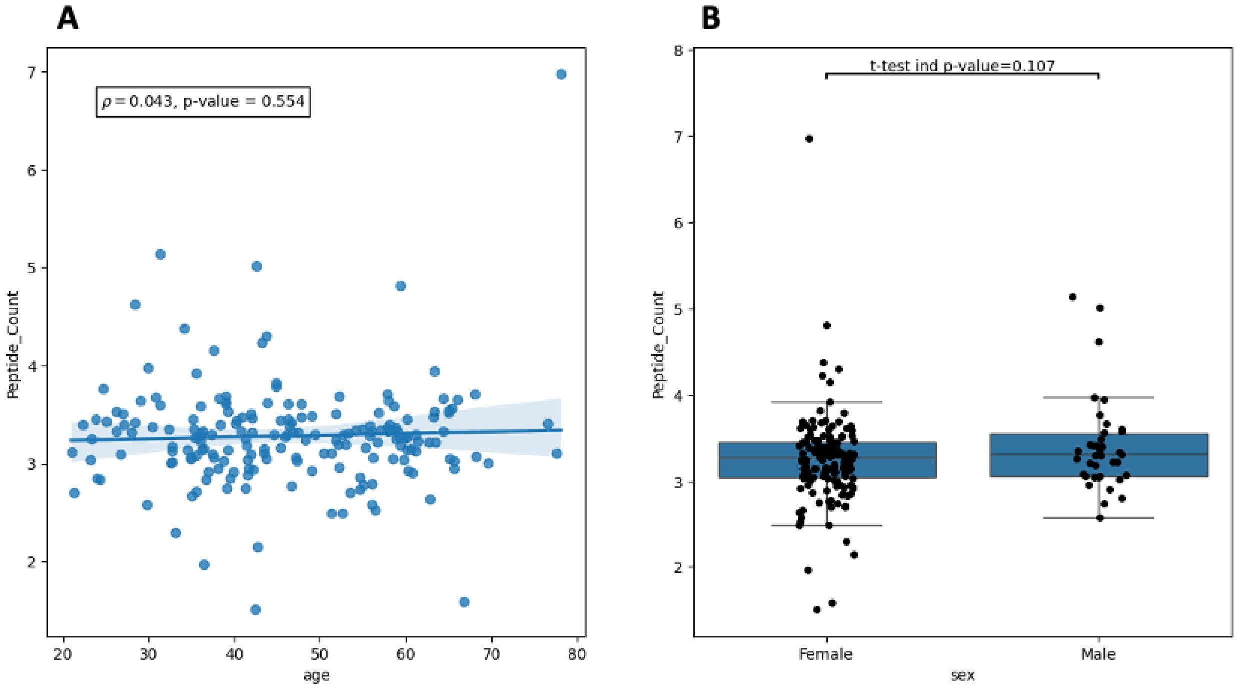


**Figure S1**. Data showing detailed plots of the peptide counts (per 1 x $10^6$) and their association with (**A**) age and (**B**) sex. Correlation with age was determined by calculating the Pearson correlation coefficient between the peptide counts and age (years). We did not observe any significant correlation with age (Pearson's R = 0.043, P-value = 0.554). Sex-specific associations were calculated via an independent T-test between the peptide counts and sex (female and male) and we did not observe any significant association (P-value = 0.107).

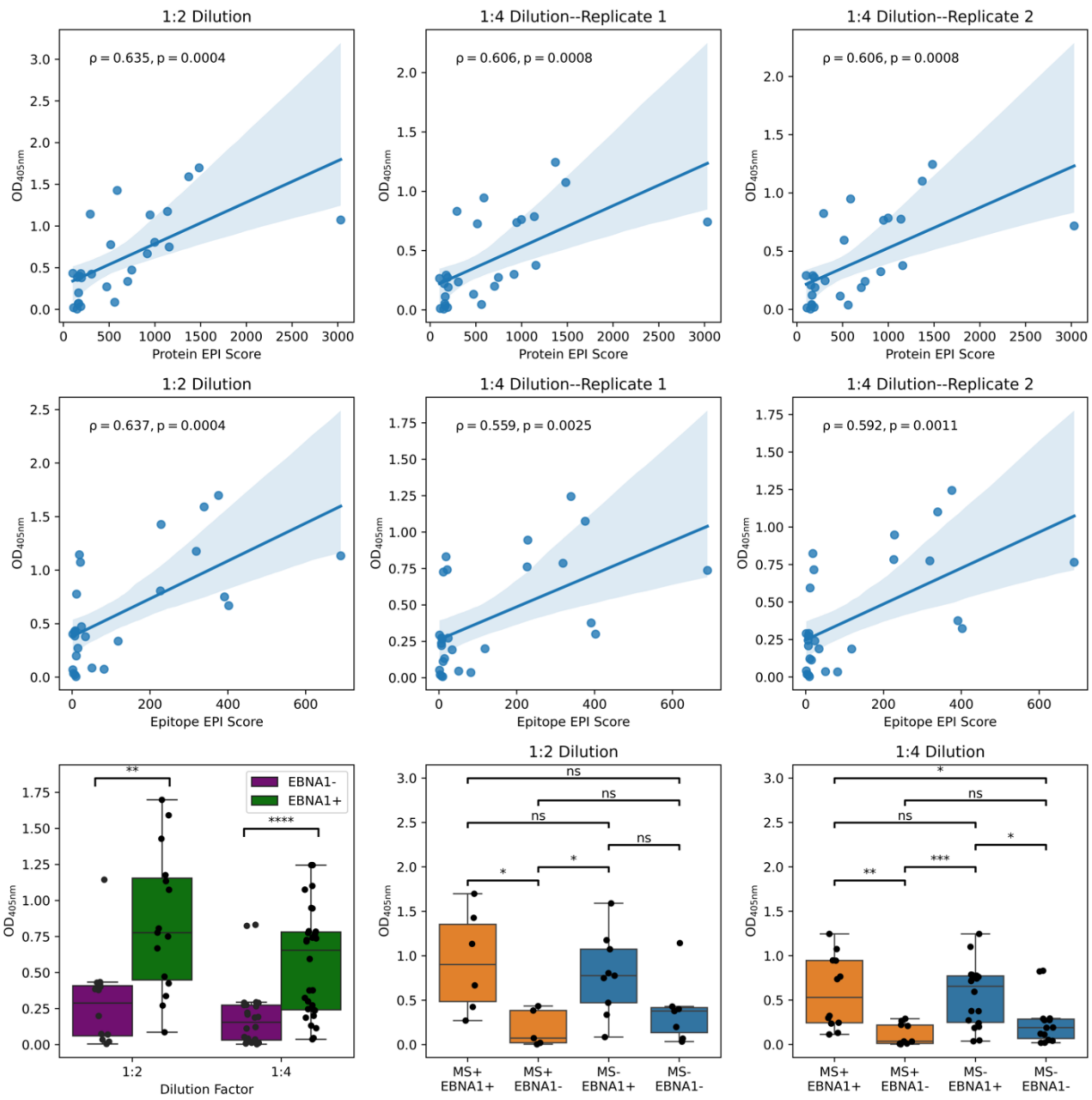


**Figure S2**. Data showing detailed plots of the ELISA assays for EBNA1+ samples versus EBNA1- samples.

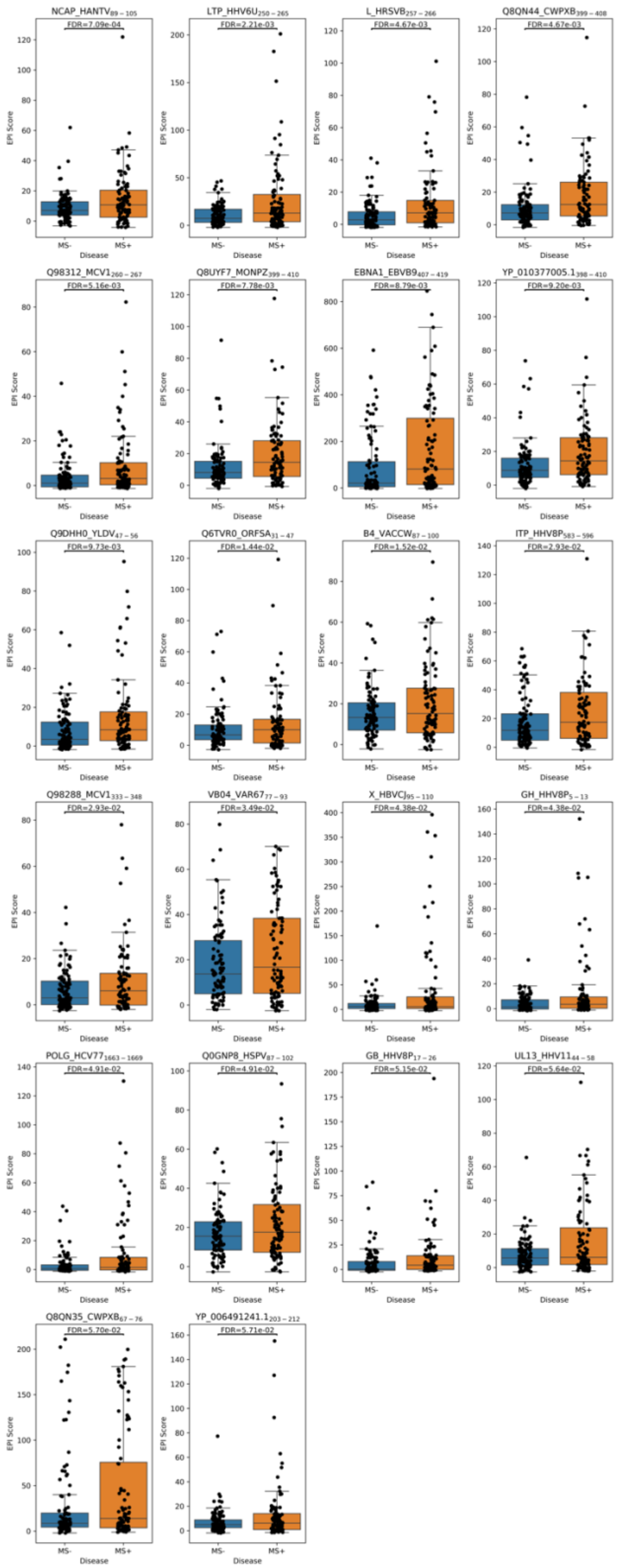


**Figure S3**. Boxplots showing the distributions of the epitope intensity score (EPI Score) for viral epitopes associated with MS.

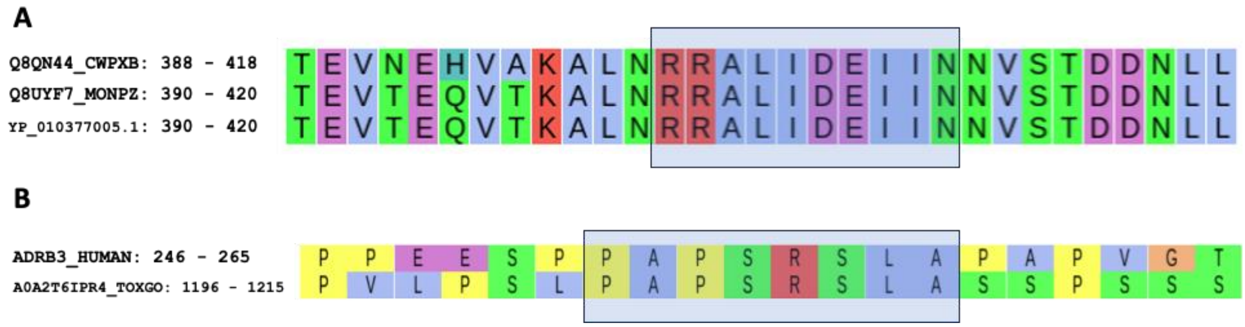


**Figure S4**. Protein sequence alignment of detected epitopes that share sequence homology. (**A**) 3 ankyrin-repeat proteins of cowpox and monkeypox virus with MS-associated epitope having shared sequence homology. (**B**) Homology between ADRB3 and the RNA polymerase Rpb1 C-terminal repeat-containing protein of Toxoplasma gondii. The boxed region highlights the associated epitope sequence shared between proteins. The alignments are performed using EMBL-EBI Muscle aligner (https://www.ebi.ac.uk/jdispatcher/msa/muscle).

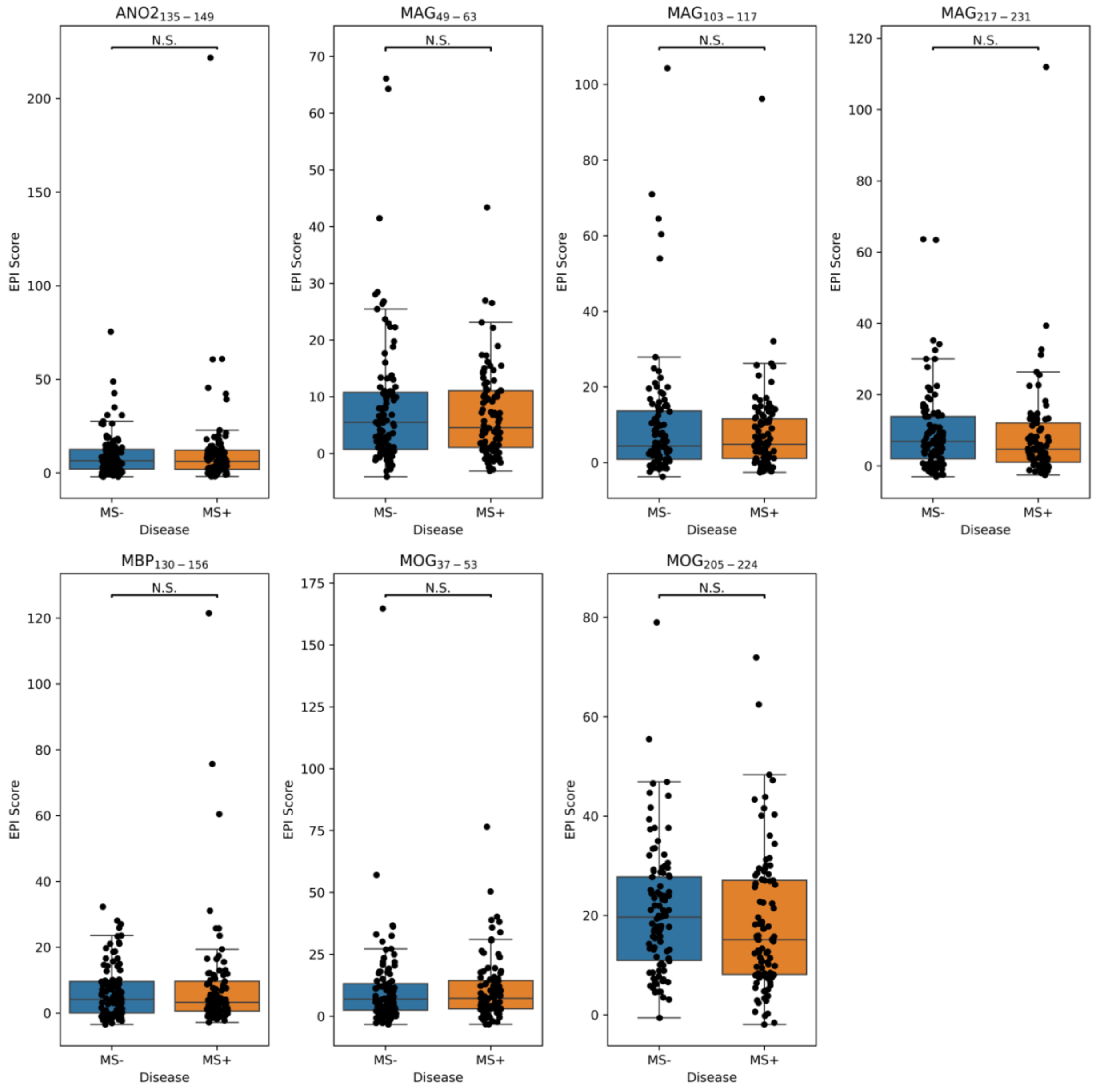


**Figure S5**. Boxplots showing the distributions of the epitope intensity score (EPI Score) for human autoantigenic epitopes identified from other studies.

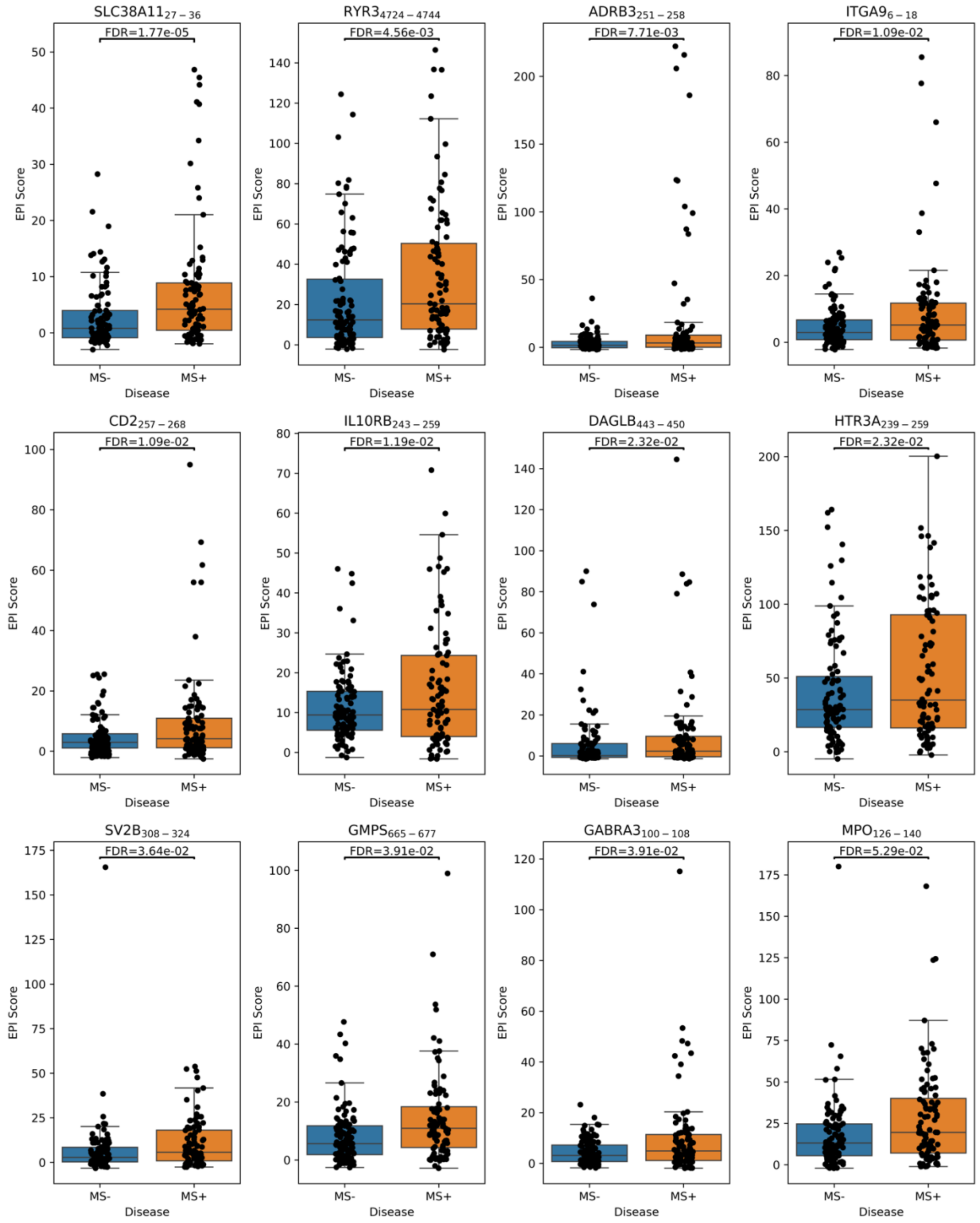


**Figure S6**. Boxplots showing the distributions of epitope intensity score (EPI Score) for human autoantigen epitopes associated with MS.

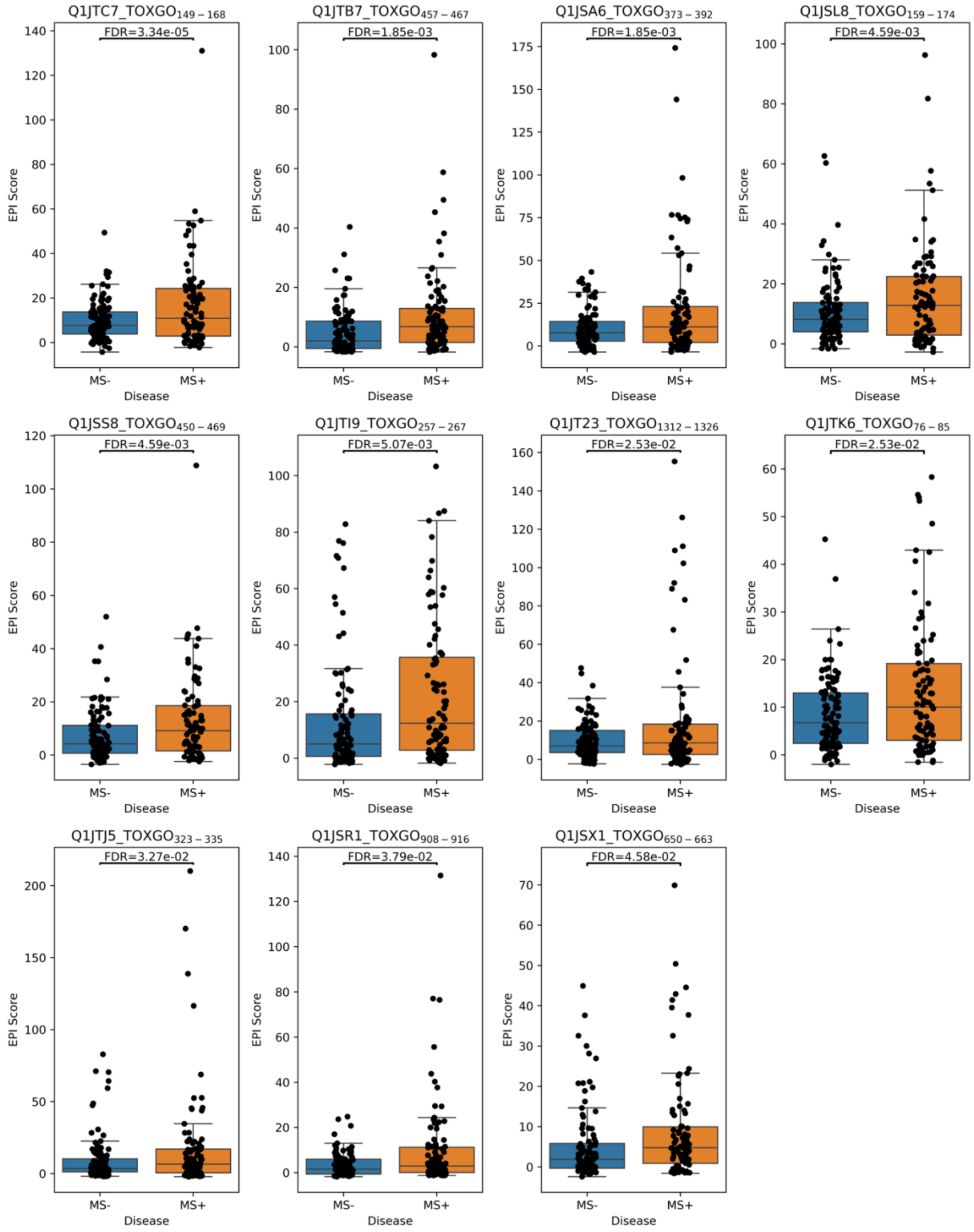


**Figure S7**. Boxplots showing the distributions of epitope intensity score (EPI Score) for Toxoplasma gondii epitopes associated with MS.

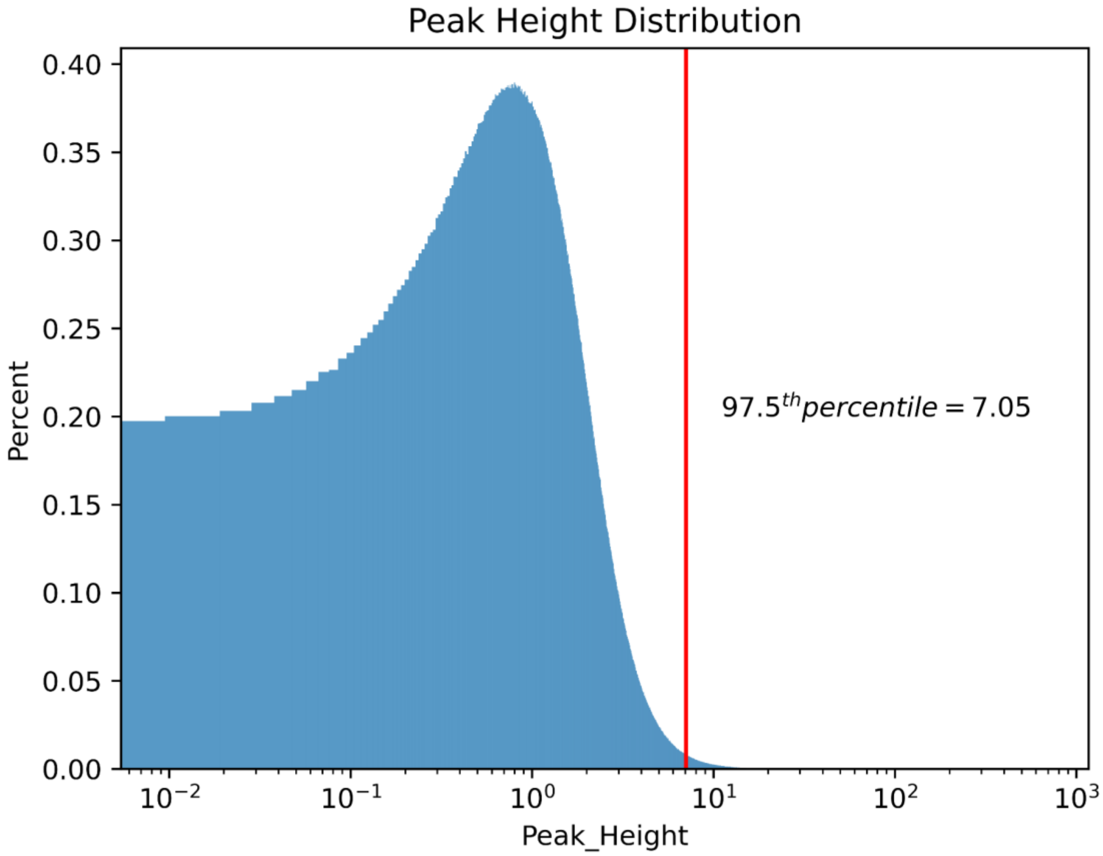


**Figure S8**. Distribution of all local maxima (peaks) heights across all samples for viral proteins. The red line represents the 97.5th percentile of peak heights. Peaks above this cutoff were considered to be a potential epitope.
